# Conformational dynamics and allosteric modulation of the SARS-CoV-2 spike

**DOI:** 10.1101/2021.10.29.466470

**Authors:** Marco A. Díaz-Salinas, Qi Li, Monir Ejemel, Leonid Yurkovetskiy, Jeremy Luban, Kuang Shen, Yang Wang, James B. Munro

**Affiliations:** Department of Microbiology and Physiological Systems, University of Massachusetts Chan Medical School, Worcester, MA 01605, USA; MassBiologics of the University of Massachusetts Chan Medical School, Boston, MA 02126, USA; Program in Molecular Medicine, University of Massachusetts Chan Medical School, Worcester, MA 01605, USA; Department of Biochemistry and Molecular Biotechnology, University of Massachusetts Chan Medical School, Worcester, MA 01605, USA

## Abstract

Severe acute respiratory syndrome coronavirus 2 (SARS-CoV-2) infects cells through binding to angiotensin-converting enzyme 2 (ACE2). This interaction is mediated by the receptor-binding domain (RBD) of the viral spike (S) glycoprotein. Structural and dynamic data have shown that S can adopt multiple conformations, which controls the exposure of the ACE2-binding site in the RBD. Here, using single-molecule Förster resonance energy transfer (smFRET) imaging we report the effects of ACE2 and antibody binding on the conformational dynamics of S from the Wuhan-1 strain and the B.1 variant (D614G). We find that D614G modulates the energetics of the RBD position in a manner similar to ACE2 binding. We also find that antibodies that target diverse epitopes, including those distal to the RBD, stabilize the RBD in a position competent for ACE2 binding. Parallel solution-based binding experiments using fluorescence correlation spectroscopy (FCS) indicate antibody-mediated enhancement of ACE2 binding. These findings inform on novel strategies for therapeutic antibody cocktails.

## Introduction

Severe acute respiratory syndrome coronavirus 2 (SARS-CoV-2) is the etiologic agent of the coronavirus disease 2019 (COVID-19) pandemic (*1*). Despite the existence of efficacious COVID-19 vaccines (*2*), urgent needs remain for preventative and therapeutic strategies to control this unprecedented situation, as well as to stop the emergence of new variants of concern (*3*).

To infect host cells, SARS-CoV-2 binds the cell receptor angiotensin-converting enzyme 2 (ACE2) through its envelope glycoprotein spike (S), which subsequently promotes membrane fusion and cell entry (*1*, *4*–*11*). S is a trimer of heterodimers, with each protomer consisting of S1 and S2 subunits (Fig. 1). S1 contains the receptor-binding domain (RBD), which includes the ACE2 receptor binding motif (RBM). S2, which forms the spike stalk, undergoes a large-scale refolding during promotion of membrane fusion (*12*–*15*). Structures of the soluble trimeric ectodomain of the SARS-CoV-2 S glycoprotein in two prefusion conformations have been reported (*10*, *11*, *16*). These distinct conformations demonstrate that the RBD of each protomer can independently adopt either a “down” (closed) or an “up” (open) position, giving rise to asymmetric trimer configurations (Fig. 1A). The RBM is occluded in the down conformation, suggesting that the RBD transitioning from the down to the up conformation is required for binding the ACE2 receptor. Indeed, structures of S bound to ACE2 show the RBD in the up conformation (*17*). Structural data were corroborated by real-time analysis of the conformational dynamics of S through single-molecule Förster resonance energy transfer (smFRET) imaging (*18*).

**Fig. 1.**
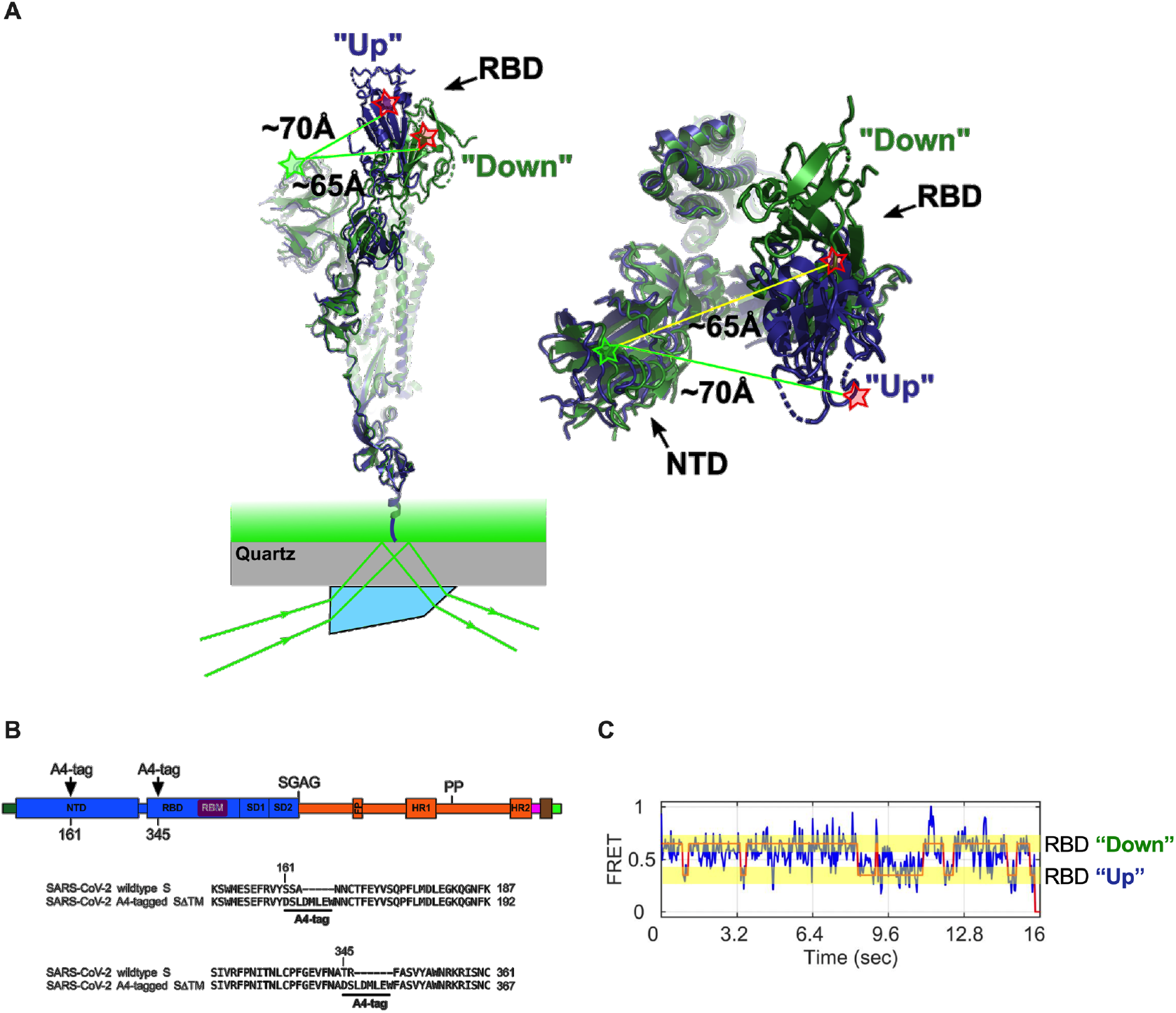
smFRET imaging the conformational dynamics of the SARS-CoV-2 S ectodomain. (**A**) (left) SARS-CoV-2 SΔTM containing a single fluorescently labelled A4-tagged protomer within an otherwise untagged trimer was immobilized on a streptavidin-coated quartz microscope slide by way of a C-terminal 8X His tag and biotin-NiNTA. For clarity, only a monomer is depicted. Individual SΔTM trimers were visualized with prism-based TIRF microscopy using a 532-nm laser. Overlay of two S protomers with RBD domains in the “up” (blue) and “down” (green) conformations are shown with approximate positions of fluorophores indicated by green (LD550) and red (LD650) stars. (right) Top view of the same S protomer overlay. The approximate distances between the sites of labeling are shown. Structures adapted from PDB 6VSB. (**B**) (top) Domain organization of the SARS-CoV-2 SΔTM construct used for smFRET experiments, indicating the sites of A4 tag insertion. The S1 and S2 subunits are in blue and orange, respectively. Additional domains and features are as follows, ordered from N- to C-terminus: signal peptide, dark green; NTD, N-terminal domain; RBD and RBM, receptor binding domain and motif (purple), respectively; SD1, subunit domain 1; SD2, subunit domain 2; SGAG, furin cleavage site mutation; FP, fusion peptide; HR1 and HR2, heptad repeat 1 and 2, respectively; PP, diproline mutations; T4 fibritin trimerization motif (foldon), magenta; TEV protease cleavage site, brown; 8X-His tag, green; SGAG, sequence replacing furin-cleavage site. (bottom) Amino acid sequence alignments indicating sites of A4-tag insertions in SΔTM. A4 peptide sequences (DSLDMLEW) are underlined. (**C**) Representative smFRET trajectory acquired from an individual SΔTM trimer (blue). Idealization resulting from HMM analysis is overlaid (red). High and low FRET states corresponding to the “down” and “up” positions of the RBD, respectively, are indicated and highlighted in transparent yellow bars.

These structural and biophysical data suggest that modulating the conformational equilibrium of the RBD of S might be a determinant of SARS-CoV-2 infectivity and neutralization sensitivity. By the summer of 2020, the SARS-CoV-2 S variant D614G (strain B.1) had supplanted the ancestral virus (strain Wuhan-1) worldwide, and structural analysis showed that D614G disrupts an interprotomer contact (*19*). This disruption results in a shift in the RBD conformation toward the up position, which is competent for ACE2 binding, consistent with increases in *in vitro* virus-cell binding mediated by ACE2 and infectivity (*16*, *20*). At the same time, the enhanced exposure of the RBM in the D614G variant led to increased sensitivity to neutralizing antibodies (*21*). Furthermore, the RBD showed stabilization in the up position, as well as an intermediate conformation, upon treatment with a neutralizing S2 stalk-directed antibody (*22*, *23*).

Here we report on the conformational dynamics of SARS-CoV-2 S in the absence or presence of ligands visualized using an smFRET imaging assay (Fig. 1A). Our results indicate that ACE2 binding is controlled by the intrinsic conformational dynamics of the RBD, with ACE2 capturing the intrinsically accessible up conformation rather than inducing a conformational change. We find that antibodies that target diverse epitopes—including epitopes in the NTD and in the S2 stalk, which are distal to the RBD—tend to shift the RBD equilibrium on the D614 spike toward the up conformation, enhancing ACE2 binding. The D614G spike existed in an equilibrium where the RBD favors the up conformation prior to antibody binding. Nonetheless, antibodies that target the S2 stalk further promoted the RBD-up conformation on the D614G spike. We thus observe long-range allosteric modulation of the RBD equilibrium, which in turn regulates exposure of the ACE2-binding site. Inducing exposure of key neutralizing epitopes with antibodies will inform the design of novel therapeutic cocktails (*24*–*26*).

## Results

### Tagged SARS-CoV-2 S spike maintains a native conformation

With the aim of visualizing the conformational dynamics in real-time of SARS-CoV-2 S, we developed an smFRET imaging assay. We specifically sought to probe the movement of the RBD between the up and down positions. To this end, guided by the available structural data (*10*, *11*) (Fig. 1A), we inserted the 8-amino-acid A4 peptide into the spike ectodomain (SΔTM) within loops located between the β7-β8 strands in the NTD at position 161, and between helix α1 and strand β1 in the RBD at position 345 (Fig. 1B). Fluorophores were then enzymatically attached to the A4 tags through incubation with AcpS (*27*). This approach was chosen because it was previously used for conformational dynamics studies of S, as well as the spike proteins from HIV-1 and Ebola (*18*, *28*–*30*). Structural analysis indicated an increase in the distance between the attachment sites of LD550 and LD650 fluorophores after the RBD transitions from the down to the up conformation, suggesting that this labeling strategy would allow us to visualize this dynamic event (Fig. 1A) (*18*).

Before proceeding to smFRET imaging, we first sought to validate the structure and antigenicity of the 161/345A4-tagged SΔTM trimer. Homo-trimers with either D614 or D614G were validated through two different approaches: (1) evaluation of their binding to ACE2 and, (2) evaluation of their antigenic characteristics compared with untagged SΔTM (Fig. S1A-B). We developed a fluorescence correlation spectroscopy (FCS) assay to evaluate ACE2 binding to A4-tagged SΔTM trimers in solution (Fig. 2A). For this assay, purified ACE2 (Fig. S1C) was conjugated to the Cy5 fluorophore (Fig. S1D). Cy5-ACE2 was incubated with varying concentrations of either tagged or untagged SΔTM and the timescale of diffusion was measured using FCS. The FCS data were fit to a model of two diffusing species (*31*) (Fig. 2B). Fitting led to determination of diffusion times for unbound ACE2 (t_free_ = 0.48±0.02 ms), and ACE2 bound to SΔTM D614 (t_D614-bound_ = 4.53±0.11 ms) or to the D614G variant (t_D614G-bound_ = 4.32±0.15 ms). As expected, the diffusion times for the SΔTM-ACE2 complex were higher than for unbound ACE2, consistent with the formation of a larger complex with slower diffusion. Moreover, FCS experiments allowed us to calculate dissociation constants (*KD*) for ACE2 binding to untagged and A4-tagged SΔTM proteins in solution (Fig. 2C), which were approximately 12.4±2.7 nM and 8.3±1.2 nM, respectively, in rough agreement with values obtained through surface-based assays (*10*, *16*). The antigenicity of A4-tagged SΔTM homo-trimers was evaluated through an ELISA assay described in Material and Methods, using the RBD-targeting antibodies MAb362 (both IgG1 and IgA1) (*32*), REGN1098 (*33*), S309 (*34*) and CR3022 (*35*); NTD-targeting antibody 4A8 (*36*), as well as the stalk-targeting antibodies 1A9 (*37*) and 2G12 (*38*) (Fig. S2A). A4-tagged SΔTM homo-trimers maintained more than 50% of antibody binding compared to untagged SΔTM (Fig. S2B), with MAb362-IgG1 and 4A8 showing no significant loss of binding. Taken together, these results suggest that double 161/345 A4-labeled SΔTM trimers maintain native functionality during ACE2 binding and near-native antigenic properties.

**Fig. 2.**
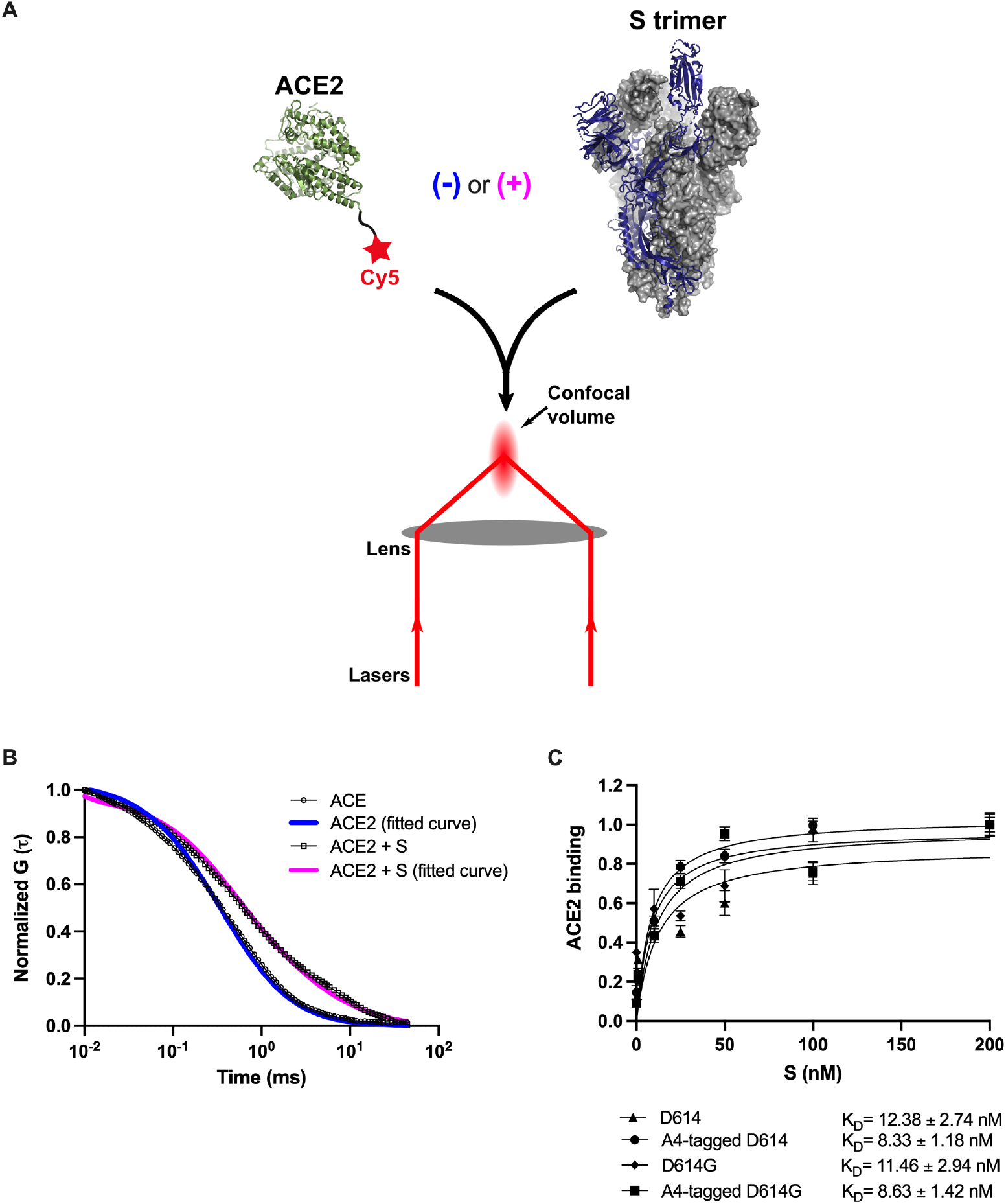
Verification of ACE2-binding to A4-tagged SΔTM trimers using FCS. (**A**) Cy5-labeled ACE2 was incubated in the absence or presence of untagged or A4-tagged SΔTM spikes. The diffusion of Cy5-ACE2 was evaluated by FCS using a 647-nm laser as indicated at Materials and Methods. (**B**) Representative normalized autocorrelation curves for Cy5-ACE2 in the absence (circles) or presence (squares) of SΔTM, and the corresponding fits are shown in blue or magenta, respectively. The shift in the autocorrelation to longer timescales seen in the presence of SΔTM reflects the slower diffusion resulting from the larger size of the complex. (**C**) Cy5-ACE2 (100 nM) was incubated with different concentrations of the indicated SΔTM spikes and the resulting mixture was evaluated by FCS as described in Material and Methods. Dissociation constants (*KD*) determined from fitting the titration are indicated for the different SΔTM constructs. Data are presented as the mean ± standard deviation from three independent measurements.

### Effects of ACE2 on the SARS-CoV-2 S RBD conformational equilibrium

To monitor the conformational dynamics of SΔTM D614 and D614G, we purified SΔTM hetero-trimers (Fig. S1E-G), formed by co-transfection of 161/345A4-tagged and untagged SΔTM plasmids at a 1:2 ratio (Material and Methods). This ensured that on average the SΔTM trimers were comprised of one tagged protomer and two untagged protomers. The SΔTM hetero-trimers were then labeled with equimolar concentrations of LD550 and LD650 fluorophores. The labeled trimers were then incubated in the absence or presence of ACE2 before immobilization on a quartz microscope slide and imaging with TIRF microscopy. smFRET trajectories acquired from individual unbound SΔTM D614 molecules showed transitions between high (~0.65) and low (~0.35) FRET states, suggestive of the down and up RBD positions, respectively (Fig. 3A). Hidden Markov modeling (HMM) confirmed that a 2-state kinetic model was sufficient to describe the dynamics observed in the smFRET trajectories. Consistent with SΔTM D614 preferring the down conformation, HMM analysis indicated 61.0±1.7% occupancy in the high-FRET state and 39.0±1.7% occupancy in the low-FRET state. The same FRET states were detected after incubation with ACE2, but the conformational equilibrium shifted to 36.8±2.1% in the high-FRET state and 63.2±2.1% occupancy in the low-FRET state (Fig. 3B–C). This result is consistent with ACE2 promoting the RBD-up conformation. HMM analysis also indicated a reduction in the overall dynamics upon ACE2 binding, as indicated by the transition density plots (TDPs; Fig. 3A–B), which display the relative frequencies of transitions between the high- and low-FRET states. The rates of transition were determined through maximum likelihood estimation. This analysis indicated that transition from the high- to the low-FRET state occurred at *k*_−1_=2.6±0.2 sec^−1^, whereas the low- to high-FRET transition occurred at *k*_1_=3.8±0.2 sec^−1^. ACE2 binding had minimal effect on the high- to low-FRET transition (*k*_−1_=2.2±0.2 sec^−1^), but reduced the low- to high-FRET transition to *k*_1_=1.3±0.1 sec^−1^ (Fig. 3D). This analysis thus specified that the effect of ACE2 binding is to capture and stabilize the up conformation (low-FRET state) and reduce transitions to the down conformation (high-FRET state). ACE2 binding does not significantly affect the stability of the down conformation, nor induce transitions to the up conformation.

**Fig. 3.**
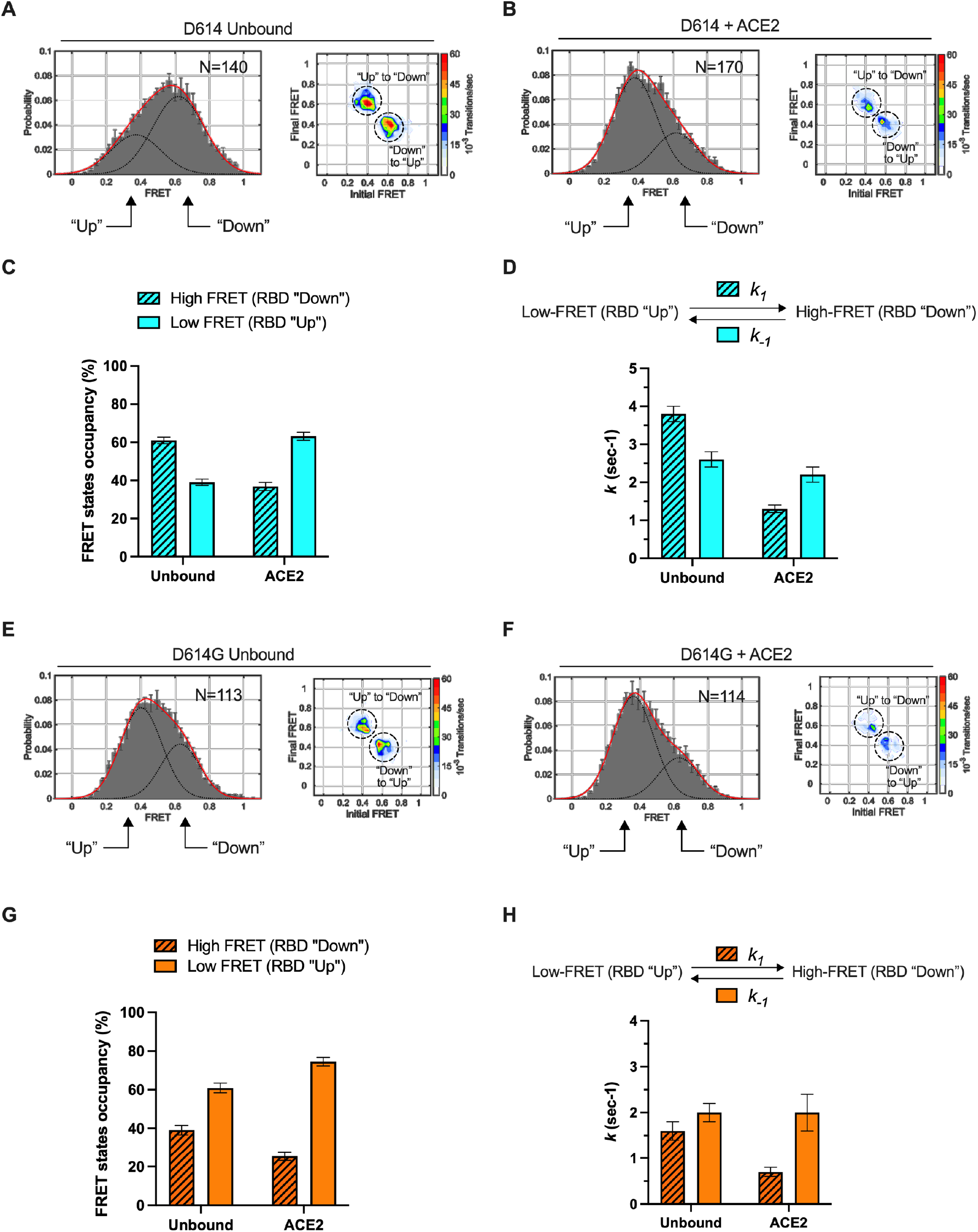
ACE2-binding modulates the RBD conformation of SΔTM D614 and D614G. (**A**) (left) FRET histogram for unbound SΔTM D614 overlaid with the sum of two Gaussian distributions (sum, red; single distributions, grey) reflecting the results of HMM analysis. The Gaussian distributions are centered at 0.65 and 0.35 FRET, corresponding to the RBD-down and RBD-up conformations, respectively. FRET histograms are presented as the mean ± standard error determined from three independent populations of smFRET traces. The number of smFRET traces compiled in the histogram is shown (N). (right) Transition density plot (TDP) for unbound SΔTM D614 indicating the frequency of observed FRET transitions, as indicated, determined through HMM analysis. (**B**) The same data for the ACE2-bound SΔTM D614 spike. (**C**) Quantification of the high- and low-FRET state occupancies for unbound and ACE2-bound SΔTM D614 determined from HMM analysis of the individual smFRET traces. Occupancies are presented as the mean ± standard deviation across the population of traces. Numeric data are shown in Table S2. (**D**) (top) Kinetic scheme defining the rates of transition between FRET states. (bottom) Rates of transition for unbound and ACE2-bound SΔTM D614 determined from HMM analysis of the individual smFRET traces. Rate constants are presented with error bars estimated from 1000 bootstrap samples. Numeric data are shown in Table S2. (**E,F**) FRET histogram and TDP for the unbound and ACE2-bound SΔTM D614G spike, displayed as in (A). (**G,H**) FRET state occupancies and rates constants for the unbound and ACE2-bound SΔTM D614G spike, displayed as in (C) and (D). Numeric data are shown in Table S3.

We next sought to determine the effect of the D614G mutation on the conformational dynamics of SΔTM. We observed the same two FRET states for SΔTM D614G as for the ancestral D614 spike (Fig. 3E). However, the unbound SΔTM D614G displayed greater occupancy in the low-FRET state (60.9±2.5%), and the overall level of dynamics was reduced as compared to D614 (Fig. 3E–G). The rate constants, *k*_−1_ and *k*_1_, were reduced to 2.0±0.2 sec^−1^ and 1.6±0.2 sec^−1^, respectively (Fig. 3H). ACE2 binding further increased the low-FRET occupancy to 74.5±2.2% and reduced the overall level of dynamics shown in the TDPs (Fig. 3F). As seen for D614, ACE2 binding had minimal effect on the rate of transition from the high- to the low-FRET state (*k*_−1_=2.0±0.4 sec^−1^) but reduced the rate of transition from the low- to the high-FRET state to *k*_1_=0.7±0.1 sec^−1^ (Fig. 3H). Thus, consistent with structural studies (*16*, *19*), the D614G mutation shifted the conformational equilibrium in favor of the RBD-up conformation. Also, here again, ACE2 binding stabilized the RBD-up conformation without affecting the energetics of the RBD-down conformation.

### RBD-targeting antibodies promote the RBD-up conformation of S D614

Numerous neutralizing monoclonal antibodies (mAbs) targeting SARS-CoV-2 S have been identified (*39*). However, their mechanisms of action have only been partially described, especially for mAbs that target epitopes outside of the RBD. We first sought to use our smFRET imaging approach to explore the effect of RBD-directed mAbs on SΔTM dynamics for both the D614 and D614G variants. We chose neutralizing RBD-directed mAbs from different classified groups according to the epitope targeted (*40*): (1) MAb362 (isoforms IgG1 and IgA1) that directly targets the RBM(*32*); (2) REGN10987, which binds an epitope located on the side of the RBD, blocking ACE2 binding without directly interacting with the RBM (*33*); and S309 and CR3022 that bind the RBD but do not compete with ACE2 binding (*34*, *35*). Imaging of SΔTM D614 pre-incubated with each of the above mAbs revealed a predominant low-FRET state associated with the RBD in the up conformation (Fig. S3, left, and Table S2). In all cases, the mAbs stabilized the low-FRET state as compared to the unbound SΔTM, although none to the extent seen for ACE2 (Fig. 4A). As observed during ACE2 binding, the mAbs generally induced a larger effect on the rate of transition from the low- to the high-FRET state (*k*_1_), with a minor effect on the high- to low-FRET transition (*k*_−1_; Fig. 4B). In contrast, none of the mAbs stabilized the low-FRET state for SΔTM D614G to a significant extent (Figs. 4A, S3 and Table S3), suggesting that the effect of the D614G mutation is sufficient to enable mAb binding without further conformational changes.

**Fig. 4.**
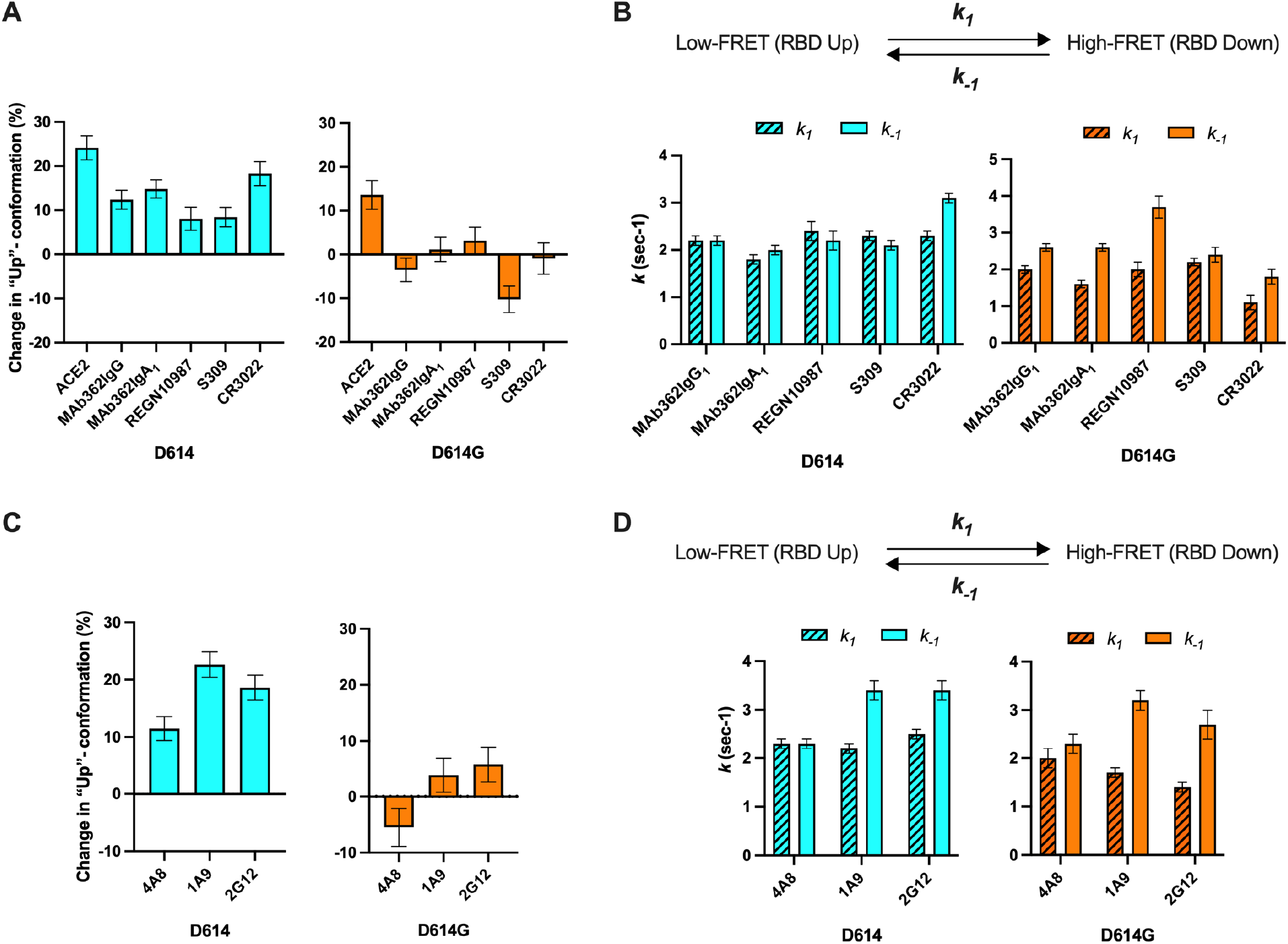
Antibodies directly and allosterically modulate SΔTM RBD conformation. (**A**) The change in low-FRET occupancy (RBD-up conformation), calculated by subtracting the low-FRET occupancy for unbound SΔTM from that seen for SΔTM in the presence of the indicated ligand (ACE2 or RBD-targeted mAb). The changes in occupancy are shown for SΔTM D614 (cyan) and D614G (orange). (**B**) (top) Kinetic scheme defining the rates of transition between FRET states. (bottom) Rates of transition for SΔTM D614 (cyan) and D614G (orange) in the presence of RBD-targeted mAbs determined from HMM analysis of the individual smFRET traces. Rate constants are presented with error bars estimated from 1000 bootstrap samples. Numeric data are shown in Tables S2 and S3. (C) The change in low-FRET occupancy (RBD-up conformation) seen for SΔTM D614 (cyan) and D614G (orange) in the presence of NTD-(4A8) and stalk-targeted (1A9 and 2G12) mAbs. Data were determined as in (A). (**D**) Rates of transition between FRET states for SΔTM D614 (cyan) and D614G (orange) in the presence of NTD and stalk-targeted mAbs, presented as in (B). Numeric data are shown in Tables S2 and S3.

### NTD- and stalk-targeting mAbs allosterically modulate the RBD position

Several mAbs have been identified that target epitopes outside of the RBD. Some of which bind the NTD and are potently neutralizing (*36*, *41*–*43*). We therefore explored the conformational dynamics of both SΔTM D614 and D614G pre-treated with the NTD-targeting mAb 4A8 (*36*), and with the S2 stalk-directed mAbs 1A9 (*37*) and 2G12 (*38*). 4A8 treatment of SΔTM D614 stabilized the low-FRET state to a comparable extent as seen for the RBD-targeted mAbs (Figs. 4C, S3, and Table S2). The change in transition rates also followed a similar trend as seen for RBD-targeted mAbs with the low- to high-FRET (*k*_1_) transition being reduced, with a minor effect on the high- to low-FRET transition (*k*_−1_; Figs. 4D, S3, and Table S2). The stalk-directed mAbs 1A9 and 2G12 also stabilized the low-FRET state (Figs. 4C, S3, and Table S2), although kinetic analysis revealed a modulation of the dynamics that was distinct from the S1-targeted mAbs. Here, the rates of low- to high-FRET transition were reduced, while the rates of high- to low-FRET transition were increased (Fig. 4D). For SΔTM D614G, 4A8 had only a minor effect on low-FRET stability or kinetics, again suggesting that the mAb binds without affecting the conformational equilibrium. However, the stalk-targeting 1A9 and 2G12 mAbs stabilized low FRET and induced increases in the rates of transition out of high FRET (Fig. 4A, S3, and Table S3). These data indicate that the S2 stalk-targeting mAbs studied here allosterically induce transition of the RDB to the up conformation on both the D614 and D614G spikes. In contrast, the RBD- and NTD-targeting mAbs studied here capture the up conformation without actively inducing a conformational change, similar to the effects of ACE2 on the RBD conformation.

### Stalk-targeting mAbs allosterically enhance ACE2 binding

We next asked if stabilization of the RBD-up conformation by NTD- and stalk-targeted mAbs would increase ACE2 binding. We therefore applied our FCS assay for ACE2 binding after pre-treating SΔTM D614 or D614G with mAbs (Material and Methods). MAb362IgA_1_ and REGN10987 mAbs were used as controls because of their documented ACE2-blocking properties (*32*, *33*). Incubation of SΔTM D614 or D614G with MAb362IgA_1_ or REGN10987 resulted in statistically significant reductions in ACE2 binding that are consistent with previous reports at comparable mAb concentrations (*32*, *33*) (Fig. 5). Overall, mAbs that stabilized the up conformation without blocking the ACE2-binding site tended to promote ACE2 binding (Fig. 5A–B). Calculation of the Spearman coefficient indicated a strong correlation (*r*_*s*_ = 0.7619) between ACE2 binding and modulation of the SΔTM RBD conformational equilibrium across all the mAbs under consideration (Fig. 5C). S309 and 4A8 provided a slight enhancement of ACE2 binding to SΔTM D614, consistent with their impacts on RBD conformation. In contrast, S309 had no significant effect on ACE2 binding to SΔTM D614G, and 4A8 had a slight inhibition of ACE2 binding, again consistent with their modulation of RBD conformation. Of particular note, the stalk-targeting 1A9 and 2G12 mAbs induced the greatest enhancement of ACE2 binding to SΔTM D614 and D614G, consistent with their allosteric modulation of RBD conformation.

**Fig. 5.**
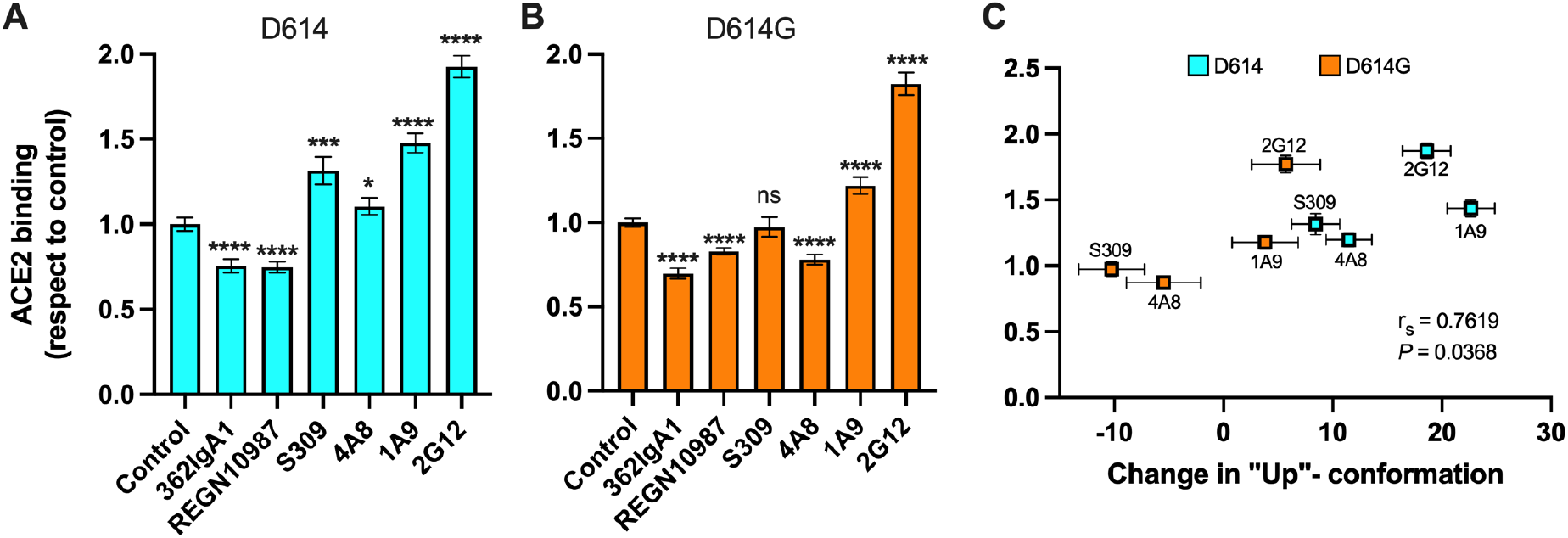
Allosteric modulation of the RBD position promotes ACE2 binding. (**A**) Binding of ACE2 by (**A**) SΔTM D614 or (**B**) D614G spikes pre-incubated with the indicated mAbs was measured by FCS as described in Material and Methods. Data are presented as the average of two independent experiments, each consisting of forty 10-sec acquisitions. Statistical significance was evaluated through a two-tailed, unpaired Mann-Whitney test as indicated in Material and Methods. *P* values <0.05 were considered significant and significance values are indicated as \**P*<0.05, \*\**P*<0.01, \*\*\**P*<0.001, \*\*\*\**P*<0.0001. (**C**) The change in the RBD-up conformation of SΔTM spikes pre-incubated with the indicated mAbs (Fig. 4A,C) exhibited a positive correlation with the binding of ACE2 determined through FCS. Statistical significance (*P*=0.0368) was found when Spearman test was performed with 95% of confidence (α=0.05).

## Discussion

Time-resolved analysis of viral spike protein conformation at single-molecule resolution complements structural studies by specifying the effects of ligand binding on the energetics of conformational dynamics. These analyses provide mechanistic insights unattainable from structures and bulk functional data alone. Here, we have developed and applied an smFRET imaging approach to monitor conformational dynamics of SARS-CoV-2 S, from the ancestral Wuhan-1 strain with D614 and the B.1 variant with D614G, during engagement with the ACE2 receptor and mAbs. Our analysis of S conformational dynamics shows that ACE2 stabilizes the RBD in the up conformation, which, in agreement with structural data, is a conformation that pre-exists prior to ACE2 binding (*10*, *11*). Determination of the kinetics of conformational changes through HMM indicated that ACE2 binding does not affect the rate of transition to the up conformation. Instead, ACE2 captures the up conformation and reduces the rate of transition to the down conformation. This can be explained by a thermodynamic stabilization of the RBD-up conformation without affecting the energetics of the down conformation (Fig. 6A). This analysis of S dynamics specifies that ACE2 binding to S does not induce a conformational change in S, but rather occurs through the capture of a pre-existing conformation.

**Fig. 6.**
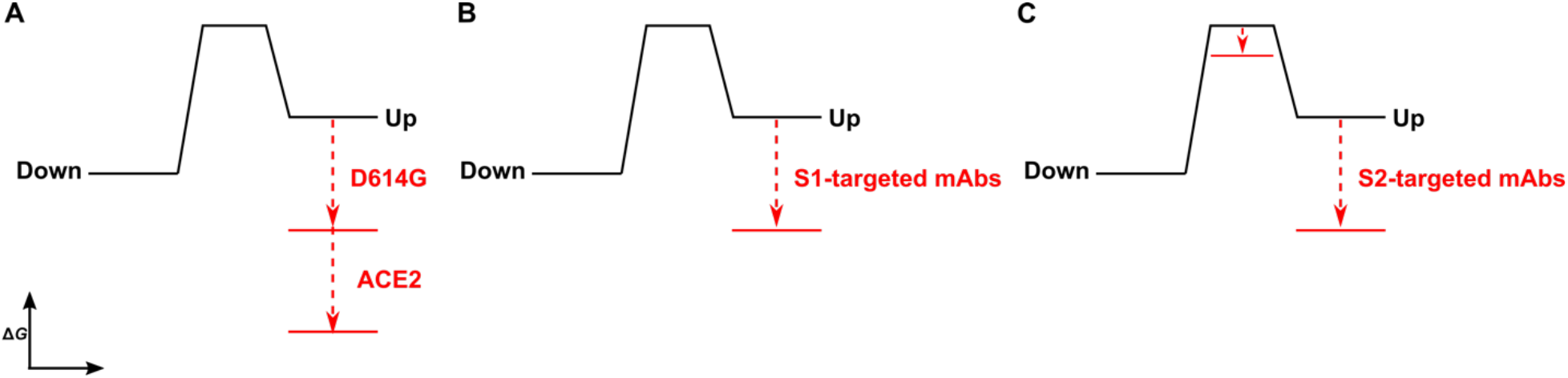
The D614G mutation and ligands modulate the S energetic landscape. (A) The D614G mutation and ACE2 have additive effects on the thermodynamic stabilization of the RBD-up conformation. (B) The predominant effect of mAbs that target the S1 domain, either the RBD (MAb362, REGN10987, S309, CR3022) or NTD (4A8), is to stabilize the RBD-up conformation. (C) mAbs that target the S2 domain have a more complex allosteric effect, resulting in stabilization of the RBD-up conformation coupled to reduction in the activation energy for transition from the RBD-down to the -up conformation.

As ACE2 binding is an essential step during SARS-CoV-2 entry, our interpretation implicates the intrinsic dynamics of S in controlling the rate or efficiency of membrane fusion during virus entry. Current models of coronaviral membrane fusion depict the RBD-up conformation as an intermediate state that is on-pathway to the post-fusion S conformation (*44*, *45*). Accordingly, factors that stabilize the RBD-up conformation would likely increase the rate of membrane fusion. Our data demonstrate that the D614G mutation stabilized the RBD-up conformation, consistent with previous reports, which likely relates to how the mutation enhances infectivity (*16*, *19*). Previous studies have shown that D614G does not increase the rate of ACE2 binding to S (*16*, *19*), as might be expected for a conformational capture binding mechanism. This may indicate that further rearrangements in the RBD, perhaps localized in the RBM, are necessary to fully engage ACE2 beyond transition to the up conformation. Analysis of the dynamics of the unbound D614G variant showed an overall reduction in dynamics as compared to D614, consistent with the increased thermostability of the S trimer with the D614G mutation (*19*). Like ACE2 binding to the D614 spike, the predominant effect of the D614G mutation was the reduction of the rate of transition to the down conformation. This reduction in the rate constant for the RBD-up to -down transition indicates an increase in the activation energy, which is mainly explained by an increase in thermodynamic stability of the RBD-up conformation (Fig. 6A). ACE2 binding to S D614G had an additive effect on the RBD position, pushing the equilibrium further toward the up conformation than either ACE2 binding or the D614G mutation did independently. Thus, the D614G mutation permits further stabilization of an intermediate conformation captured by ACE2 binding. Here again, ACE2 binding functioned by specifically increasing the thermodynamic stability of the RBD-up conformation. Residue D614 is distal to the RBD and forms a salt bridge with K854 in the fusion-peptide proximal region, which is lost with the D614G mutation (*17*, *19*, *46*). Our analysis shows that the D614-K854 electrostatic interaction had a destabilizing effect on the thermodynamics of the RBD-up conformation. The similar impacts of D614G and ACE2 binding on the S energetic landscape implies that the mutation provides a fitness advantage by mimicking the effects of receptor binding. Such a long-range allosteric connection between the receptor-binding domain and the region surrounding the fusion peptide has been reported for the influenza hemagglutinin and the Ebola virus envelope glycoprotein (*47*–*49*), suggestive of a common mechanistic connection between receptor binding and triggering movement of the fusion peptide (or fusion loop) among class-I viral fusion proteins.

We find that mAbs that target S1 of the D614 spike, including the RBD and NTD, have a similar impact on conformational dynamics as ACE2, with the predominant effect being the reduction in the rate of transition to the down conformation. The overall minimal effect on the rate of transition to the up conformation is again consistent with thermodynamic stabilization and the capture of a pre-existing S conformation (Fig. 6B). mAb S309 had a notably modest effect on the stability of the up conformation, consistent with structural data demonstrating that it binds to the RBD in either the up or down positions (*34*). Overall, RBD-and NTD-targeting mAbs had minimal effect on the conformation of the D614G spike. The exception was mAb S309, which modestly destabilized the up conformation, shifting the equilibrium to approximately that seen for the D614 spike bound to S309. As S309 does not prevent ACE2 binding, its mechanism of broad neutralization remains unclear (*34*, *50*). However, its modulation of the RBD position likely plays some role and may impact downstream conformational changes related to membrane fusion.

Our kinetic analyses have shown that the S1-targeted ligands considered here capture the up conformation. In contrast, the S2-targeted mAbs considered here induce conformational changes in the RBD by reducing the activation energy for transition into the up conformation, while also stabilizing the up conformation (Fig. 6C). Cryo-electron tomography of SARS-CoV-2 virions has revealed the presence of three flexible hinges within the S stalk: the hip, knee, and ankle. These hinges connect the head, and the upper and lower legs of S and confer flexibility on the spikes (*51*, *52*). Our smFRET data demonstrate that stalk-targeted mAbs 1A9 and 2G12 allosterically modulate the position of the RBD, enhancing ACE2 binding. mAb 1A9, which neutralizes SARS-CoV, binds an epitope on S in the upper leg of the stalk near the hip and upstream of the heptad repeat helix 2 (*53*). High sequence conservation in the 1A9 epitope suggests a similar binding site in SARS-CoV-2 S and mode of action in preventing viral membrane fusion (*37*). The stalk epitope recognized by mAb 2G12, which does not neutralize SARS-CoV-2, is located near the hip and is comprised entirely of glycans (*38*) (Fig. S2A). Taken together, our data on 1A9 and 2G12 implicate the hip hinge as a critical center for allosteric control of the RBD position. Further support for the existence of allosteric centers in S2 came from other smFRET analyses of mAb CV3-25 (*54*), which binds an epitope in the upper leg of the stalk near the knee (*23*). CV3-25 was also found to promote the RBD-up conformation (*22*, *23*). Further studies are necessary to determine whether mAbs that target the lower leg and ankle hinge also exert allosteric control of the RBD.

The use of therapeutic mAb cocktails is a promising strategy, which has been explored for the treatment and prevention of Ebola virus disease (*55*). Similarly, enhancement of neutralization of SARS-CoV-2 was observed with the simultaneous use of S309 and S2E12 (*56*, *57*) which targets the RBM. This likely stems from the combined effect of S309 on S conformation and blocking ACE2 binding by S2E12. Our results suggest similar synergy in neutralization might come from the combination of 4A8 with RBM-directed mAbs. Indeed, human trials are underway evaluating mAb cocktails for COVID-19 treatment. But none of these have considered the simultaneous use of mAbs targeting the RBD and stalk of SARS-CoV-2 S (*25*, *39*). The promotion of the RBD-up conformation, which exposes the ACE2-binding site, by NTD-directed mAbs like 4A8, or stalk-directed mAbs like 1A9 and 2G12, presents a strategy for enhancement of neutralization through combination therapies with RBM-directed mAbs. The results presented here suggest the potential for synergistic inhibition of virus entry and increased potency through the combination of mAbs that target diverse epitopes.

## Materials and Methods

### Cell culture

ExpiCHO-S™ and Expi293F™ cell lines (Gibco™, Thermo Fisher Scientific, Waltham, MA, USA) were cultured in ExpiCHO™ Expression and Expi293 Expression media (Gibco™, Thermo Fisher Scientific, Waltham, MA, USA), respectively. Both cell lines were maintained at 37 °C, 8% CO_2_ with orbital shaking according to manufacturer instructions.

### Antibodies

Monoclonal antibodies MAb362 isotypes IgG1 and IgA1 has been described before(*32*). REGN10987, S309 and CR3022 antibodies heavy and light variable region sequences(*33*, *34*, *58*) were synthesized and cloned into pcDNA3.1 vector (Invitrogen™, Thermo Fisher Scientific, Waltham, MA, USA) in-frame with human IgG heavy or light chain Fc fragment. The recombinant constructs of heavy and light chain were transfected at 1:1 ratio into Expi293F™ cells using the ExpiFectamine™ 293 Transfection Kit (Gibco™, Thermo Fisher Scientific, Waltham, MA, USA). 4-5 days after transfection the antibodies were purified from the supernatant by protein A affinity resin (ProSep^®^-vA ultra, Millipore^®^, Burlington, MA, USA) and dialyzed into phosphate buffered saline pH 7.2 (PBS) overnight at 4 °C. 2G12 monoclonal antibody was expressed in ExpiCHO-S™ cells through co-transfection of plasmids encoding light and IgG heavy chains(*59*), using the ExpiFectamine™ CHO transfection kit (Gibco™, Thermo Fisher Scientific, Waltham, MA, USA) according to manufacturer instructions. The antibody was purified from the cell culture supernatant 12 days post-transfection through protein G affinity resin (Thermo Fisher Scientific, Waltham, MA, USA), and buffer exchanged and concentrated in PBS using centrifugal concentrators (Sartorius AG, Göttingen, Germany; Millipore^®^, Burlington, MA, USA). Monoclonal antibodies 4A8 and 1A9 were purchased from BioVision (Milpitas, CA, USA) and GeneTex (Irvine, CA, USA), respectively. Anti-6x-His-tag polyclonal antibody, and both HRP-conjugated anti-mouse IgG Fc and anti-human IgG Fc were purchased from Invitrogen™ (Waltham, MA, USA). Both horseradish peroxidase (HRP) conjugated anti-human kappa and anti-rabbit IgG were purchased from SouthernBiotech (Birmingham, AL, USA) and Abcam (Cambridge, UK), respectively.

### Plasmids and site-directed mutagenesis

The mammalian codon-optimized gene coding SARS-CoV-2 (Wuhan-Hu-1 strain, GenBank ID: MN908947.3) glycoprotein ectodomain (SΔTM) (residues Q14–K1211) with SGAG substitution at the furin cleavage site (R682 to R685), and proline substitutions at K986 and V987, was synthesized by GenScript^®^ (Piscataway, NJ, USA) and inserted into pcDNA3.1(−). A C-terminal T4 fibritin foldon trimerization motif, a TEV protease cleavage site, and a His-tag were synthesized downstream of the SARS-CoV-2 SΔTM (Fig. 1B). Insertion of A4 peptide (DSLDMLEW) at amino acid position 161 in SARS-CoV-2 SΔTM was done through overlap-extension PCR(*60*). Primers 2S-Age-I-F, 2S-161A4-2, 2S-161A4-3, and 2S-ApaI-R (Table S1) were used in the PCR reactions to obtain a final product bearing the 161A4 insertion, which was cloned into SΔTM using the AgeI and ApaI restriction sites. A similar strategy was performed for the A4 insertion at position 345 of SΔTM using 2S-XhoI-F, 2S-345A4-2, 2S-345A4-3, and 2S-ApaI-R primers (Table S1) in the PCR reactions. The final PCR product bearing the 345A4 insertion was cloned into SΔTM using the XhoI and ApaI restriction sites. To generate the 161/345A4 double-tagged construct, the 345A4 insertion was subcloned into the 161A4 construct through XhoI and ApaI digestion. Mutagenic PCR to obtain the D614G amino acid change into both untagged and 161/345A4-tagged SΔTM constructs was done using the primers S2_D614_Q5-F and S2_D614_Q5-R (Table S1) and the Q5^®^ Site-Directed Mutagenesis Kit (NEB^®^, Ipswich, MA, USA) according to the manufacturer instructions. Insertions and site-directed mutagenesis were confirmed through Sanger sequencing (GENEWIZ^®^, Cambridge, MA, USA).

### Protein expression and purification

Expression SΔTM trimers was performed by transfection of ExpiCHO-S™ cells with the plasmids described above using the ExpiFectamine™ CHO transfection kit (Gibco™, Thermo Fisher Scientific, Waltham, MA, USA) and according to manufacturer instructions. SΔTM hetero-trimers for smFRET experiments were expressed by co-transfection with both the untagged SΔTM (D614 or D614G) construct and the corresponding 161/345A4-tagged SΔTM plasmid at a 2:1 molar ratio. Untagged SΔTM trimers or A4-tagged hetero-trimers were purified from cell culture supernatants as follows. Supernatants containing soluble SΔTM trimers were harvested nine days post-transfection and adjusted to 20 mM imidazole, 1 mM NiSO_4_, and pH 8.0 before binding to the Ni-NTA resin. The resin was washed, and protein was eluted from the column with 300 mM imidazole, 500 mM NaCl, 20 mM Tris-HCl pH 8.0, and 10% (v/v) glycerol. Elution fractions containing SΔTM were pooled and concentrated by centrifugal concentrators (Sartorius AG, Göttingen, Germany). The SΔTM protein was then further purified by size exclusion chromatography on a Superdex 200 Increase 10/300 GL column (GE Healthcare, Chicago, IL, USA) (Fig. S1). Double 161/345A4-tagged SΔTM homo-trimers for functional assays were extracted from ExpiCHO-S™ cells at 6 days pot-transfection with a non-denaturing lysis buffer (20 mM Tris-HCl pH 8.0, 500 mM NaCl, 10% (v/v) glycerol, 1% (v/v) Triton™ X-100, 2 mg/ml aprotinin, 1 mg/ml leupeptin, and 1 mg/ml pepstatin A (Sigma-Aldrich^®^, St. Louis, MO, USA)). After 20 minutes of centrifugation at 4000 x*g*, the soluble fraction was diluted with two volumes of the same buffer without Triton™ X-100. These extracts were then passed through a 0.45 mm polyethersulfone filter unit (Nalgene™, Thermo Fisher Scientific, Waltham, MA, USA), and the tagged SΔTM was purified by affinity chromatography using Ni-NTA agarose beads (Invitrogen™, Waltham, MA, USA) and size exclusion chromatography as described above.

A plasmid encoding soluble monomeric ACE2 with a C-terminal 6x-His tag was transfected into ExpiCHO-S™ cells as described above. Supernatant containing ACE2 was harvested six days post-transfection, dialyzed at 4 °C into 20 mM Tris-HCl pH 8.0, 500 mM NaCl and 10% (v/v) glycerol buffer, using a 10 kDa MWCO dialysis membrane (Spectrum^®^ Repligen, Waltham, MA, USA). For ACE2 purification, the dialyzed supernatant was supplemented with 20 mM imidazole pH 8.0 before purification as described above for SΔTM. Purified protein concentrations were estimated by UV absorbance at 280 nm and Bradford assay (Thermo Fisher Scientific, Waltham, MA, USA). SΔTM concentration was also estimated by densitometric analysis of protein bands on immunoblots with the monoclonal antibody 1A9 as described below, and using ImageJ software v1.52q (NIH, USA).

### PAGE and immunoblots

Protein expression was evaluated by denaturing PAGE in 4-20% acrylamide (Bio-Rad, Hercules, CA, USA) and either staining with Coomassie blue or with immunoblots performed as follows. Protein gels were transferred into nitrocellulose membranes (Bio-Rad, Hercules, CA, USA) according to the manufacturer instructions. After one hour of blocking with 5% (w/v) skim milk in 0.1% (v/v) Tween™-20 (Fisher Scientific, Hampton, NH, USA) and PBS (PBS-T), membranes were incubated by shaking overnight at 4 °C with dilutions 1:2000 in blocking buffer of the primary antibody. We used a rabbit anti-6X-His antibody (Invitrogen™, Waltham, MA, USA) to detect histidine-tagged proteins or mouse 1A9 antibody (GeneTex, Irvine, CA, USA) for specific detection of SARS-CoV-2 SΔTM. Membranes were washed three times with PBS-T and then incubated with secondary HRP-conjugated anti-rabbit IgG (Abcam, Cambridge, UK) or anti-mouse IgG (Invitrogen™, Waltham, MA, USA) antibodies diluted in 0.5% (w/v) skim milk/PBS-T and incubated for one hour at room temperature. After three washes with PBS-T, membranes were developed using SuperSignal™ West Pico PLUS Chemiluminescent Substrate (Thermo Scientific™, Waltham, MA, USA) according to the manufacturer’s instructions.

### ELISA assays

96-well polystyrene plates (Thermo Scientific™, Waltham, MA, USA) were coated either with 200 ng of SARS-CoV-2 SΔTM proteins or bovine serum albumin (BSA, Thermo Scientific™, Waltham, MA, USA) through incubation overnight at 4 °C. Plates were washed three times with PBS and blocked for one hour at room temperature with the immunoblot blocking buffer described above. After three washes with PBS, plates were incubated with 600 nM of the indicated antibodies diluted in PBS for two hours at room temperature. As secondary antibodies, HRP-conjugated anti-human kappa antibody (SouthernBiotech, Birmingham, AL, USA) diluted 1:4000 in PBS was used in wells treated with MAb362, CR3022 and S309 antibodies, while HRP-conjugated anti-human IgG Fc (Invitrogen™, Waltham, MA, USA) diluted 1:10,000 in PBS was used in wells treated with REGN10987, 4A8 and 2G12 antibodies. A 1:5000 dilution of HRP-conjugated anti-mouse IgG Fc antibody in PBS was used in 1A9 antibody-treated wells.

Plates were incubated with the secondary antibody dilutions for one hour at 37 °C and developed with 1-Step™ Ultra TBM-ELISA (Thermo Scientific™, Waltham, MA, USA) reagent according to the manufacturer’s instructions. The absorbances at 450 nm were measured using a Synergy H1 microplate reader (BioTek^®^ Winooski, VT). Absorbance values from non-specific binding to BSA-coated wells were subtracted from the values obtained for SΔTM-coated wells. The background-subtracted absorbance values were then normalized to the values obtained from antibodies binding to untagged SΔTM.

### Fluorescent labeling of proteins

Purified A4-tagged SΔTM hetero-trimers for smFRET imaging were prepared by overnight incubation at room temperature with 5 μM each of coenzyme A (CoA)-conjugated LD550 and LD650 fluorophores (Lumidyne Technologies, New York, NY, USA), 10 mM MgOAc, 50 mM HEPES pH 7.5, and 5 μM Acyl carrier protein synthase (AcpS). Labeled protein was purified away from unbound dye and AcpS by size exclusion chromatography as above described, and elution fractions containing labeled SΔTM hetero-trimers were pooled and concentrated. Aliquots were stored at −80 °C until use. Purified ACE2 was labeled with Cy5 conjugated to n-hydroxysuccinimide ester (Cytiva, Marlborough, MA, USA) according to the manufacturer’s instructions. ACE2 was then purified away from unbound dye by Ni-NTA affinity chromatography as described above, followed by buffer exchanged into PBS pH 7.4 using 10 kDa MWCO centrifuge concentrators (Millipore^®^, Burlington, MA, USA).

Purified LD550/LD650-labeled SΔTM spikes and Cy5-labeled ACE2 samples were analyzed by denaturing PAGE and in-gel fluorescence was visualized using a Typhoon 9410 variable mode imager (GE Amersham Biosciences, Amersham, UK) by laser excitation at 532 nm (emission filter: 580 BP 30 Cy3) to detect LD550, or 633 nm (emission filter: 670 BP 30 Cy5) to detect LD650 or Cy5 (Fig. S1).

### smFRET imaging

Labeled SΔTM spikes (100-200 nM) were incubated in the absence or presence of unlabeled ACE2 or the indicated antibody at a monomer:ACE2 or monomer:antibody ratio of 1:3 for 90 minutes at room temperature. The 6X-His tagged SΔTM was then immobilized on streptavidin-coated quartz microscope slides by way of Ni-NTA-biotin (vendor) and imaged using wide-field prism-based TIRF microscopy as described(*28*, *29*, *62*, *63*). Imaging was performed in the continued presence of ligands at room temperature and smFRET data were collected using Micromanager(*64*) v2.0 (micro-manager.org) at 25 frames/sec. All smFRET data were processed and analyzed using the SPARTAN software (www.scottcblanchardlab.com/software) in Matlab (Mathworks, Natick, MA, USA)(*65*). smFRET traces were identified according to following criteria: mean fluorescence intensity from both donor and acceptor were greater than 50, duration of smFRET trajectory exceeded 5 frames, correlation coefficient calculated from the donor and acceptor fluorescence traces ranged between −1.1 to 0.5, and signal-to-noise ratio was greater than 8. Traces that fulfilled these criteria were then verified manually. smFRET trajectories were idealized to a 3-state hidden Markov model and the transition rates were optimized using the maximum point likelihood algorithm(*66*), implemented in SPARTAN.

### FCS-based ACE2-binding assay

ACE2 binding to the untagged and A4-tagged SΔTM spikes was evaluated by FCS as follows. Several concentrations ranging from 0.1 to 200 nM SΔTM were incubated with 100 nM Cy5-labeled ACE2 in PBS pH 7.4 for one hour at room temperature. Where indicated, 200 nM SΔTM was incubated with 600 nM of the indicated antibody for one hour at room temperature, before adding 100 nM Cy5-labeled ACE2. Non-specific antibody binding to Cy5-labeled ACE2 was determined by incubation in the absence of SΔTM. Samples were then placed on No. 1.5 coverslips (ThorLabs, Newton, NJ) and mounted on a CorTector SX100 instrument (LightEdge Technologies, Beijing, China) equipped with a 638-nm laser. 10-25 autocorrelation measurements were made for 10 sec each at room temperature for each experimental condition. To obtain the fractions of unbound and bound (*f*) ACE2 after incubation with SΔTM, normalized autocorrelation curves were fit to a model of the diffusion of two species in a three-dimensional Gaussian confocal volume(*67*, *68*),

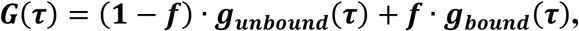

Where

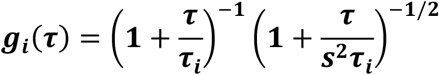

and *t*_*i*_ is the diffusion time for bound or unbound ACE2 and *s* is the structure factor that parameterizes the dimensions of the confocal volume. To determine *t*_*unbound*_ FCS data was obtained for ACE2 in the absence of SΔTM and fit to a model of a single diffusing species (*f* = 0). This value was then fixed during fitting of the FCS data obtained after incubation of ACE2 with SΔTM, so that only *t*_*bound*_ and *f* were allowed to vary. ACE2 binding was expressed as the average bound fraction (*f*) at each SΔTM concentration normalized either to the fraction bound at the highest SΔTM concentration (Fig. 2C), or to the fraction bound in the absence of antibodies (Fig. 5). All fitting was performed with a non-linear least-squares algorithm in MATLAB (The MathWorks, Waltham, MA, USA). Dissociation constants (K_D_) were determined using GraphPad Prism version 9.2.0 (GraphPad Software, San Diego, CA, USA).

### Structural analysis

Protein structures from RCSB PDB were visualized and analyzed using PyMOL™ software version 2.0.7 (The PyMOL Molecular Graphic System, Schrödinger® Inc. New York, NY, USA).

### Correlation and statistical analysis

Data sets subjected to statistical analysis were first tested for normality using GraphPad Prism version 9.2.0 (GraphPad Software, San Diego, CA, USA). Where indicated, statistical significances were evaluated through either two-tailed parametric (unpaired t-test with Welch’s correction) or nonparametric (unpaired Mann-Whitney) tests. Both tests were performed with 95% confidence levels and *P* values <0.05 were considered significant. Significance values are indicated as \**P*<0.05, \*\**P*<0.01, \*\*\**P*<0.001, \*\*\*\**P*<0.0001. Two-tailed nonparametric Spearman test with 95% confidence was performed to evaluate the correlation level between the occupancy of SΔTM in the open conformation due to allosteric antibody binding and ACE2 binding (Figs. 4 and 5). The correlation level between the above variables was determined according to established criteria(*69*) regarding Spearman coefficients (*r*_*s*_) rank values as follows: 0.00-0.10 = “negligible”, 0.10-0.39 = “weak”, 0.40-0.69 = “moderate”, 0.70-0.89 = “strong”, and 0.90-1.00 = “very strong” correlation.

## Acknowledgments

The authors thank Dr. Natasha Durham (UMass Chan Medical School) for critical discussion and reading of the manuscript.

## Funding

UMass Chan Medical School COVID-19 Pandemic Relief Fund (JBM)

National Institutes of Health grant R37AI147868 (JL)

National Institutes of Health grant R01AI148784 (JL, JBM)

Evergrande COVID-19 Response Fund (JL)

Massachusetts Consortium on Pathogen Readiness (JL)

National Institutes of Health grant K22CA241362 (KS)

Worcester Foundation for Biomedical Research (KS)

## Author contributions

Conceptualization: JBM

Methodology: MAD-S, JBM

Investigation: MAD-S, QL, ME, LK

Visualization: MAD-S, JBM

Supervision: JL, KS, YW, JBM

Writing—original draft: MAD-S

Writing—review & editing: all authors

## Data and materials availability

All data are available in the main text or the supplementary materials.

