## Supplemental Material for "Conformational dynamics and allosteric modulation of the SARS-CoV-2 spike" (from another version of this preprint: version 1, 2021-11-01)

**Supplementary Material**

**Table S1.** Primers used in this study.

| Primer name | Sequence |
|---|---|
| 2S-AgeI-F | GCT GAA ACC GGT ACC CAG TGC |
| 2S-XhoI-F[a] | GAC CTC GAG GGC AAG CAG |
| 2S-161A4-2 | CAC TCC AGC ATG TCT AGA CTG TCG TAG ACT CTG AAC TCG CTT TCC |
| 2S-161A4-3 | CAG TCT AGA CAT GCT GGA GTG GAA TAA CTG TAC ATT CGA ATA CGT GTC |
| 2S-345A4-2 | CAC TCC AGC ATG TCT AGA CTG TCG GCG TTG AAC ACC TCT CCG AAA GG |
| 2S-345A4-3 | CAG TCT AGA CAT GCT GGA GTG GTT CGC CTC TGT GTA CGC CTG GAA CAG |
| 2S-ApaI-R[a] | CTG TCA GGG CCC TAT TCA G |
| S2_D614G_Q5-F | GCTGTACCAGGGCGTGAACTGCA |
| S2_D614G_Q5-R | ACGGCCACCTGGTTGCTG |
| 2S-NdeI-F[a] | CAA GTG TAT CAT ATG CCA AGT ACG C |

[a]Primers used for Sanger sequencing in addition to cloning and mutagenesis.

**Table S2**. FRET-state occupancies and rate constants for SΔTM D614.

| SARS-CoV-2 spikes D614 | FRET-state occupancies (%) | | Rate constants ($sec^{-1}$) | |
|---|---|---|---|---|
| | Low-FRET (0.35) RBD "up"-conformation | High-FRET (0.65) RBD "down"-conformation | $k_1$ (Low to high FRET) | $k_{-1}$ (High to low FRET) |
| Unbound | 39.0±1.7 | 61.0±1.7 | 3.8±0.2 | 2.6±0.2 |
| + ACE2 | 63.2±2.1 | 36.8±2.1 | 1.3±0.1 | 2.2±0.2 |
| + Mab362IgG$_1$ | 51.4±1.3 | 48.6±1.3 | 2.2±0.1 | 2.2±0.1 |
| + Mab362IgA$_1$ | 53.9±1.2 | 46.1±1.2 | 1.8±0.1 | 2.0±0.1 |
| + REGN10987 | 47.1±2.0 | 52.9±2.0 | 2.4±0.2 | 2.2±0.2 |
| + S309 | 47.5±1.4 | 52.5±1.4 | 2.3±0.1 | 2.1±0.1 |
| + CR3022 | 57.3±2.1 | 42.7±2.1 | 2.3±0.2 | 3.1±0.2 |
| + 4A8 | 50.5±1.2 | 49.5±1.2 | 2.3±0.1 | 2.3±0.1 |
| + 1A9 | 61.7±1.4 | 38.3±1.4 | 2.2±0.1 | 3.4±0.2 |
| + 2G12 | 57.6±1.4 | 42.4±1.4 | 2.5±0.1 | 3.4±0.2 |

**Table S3**. FRET-state occupancies and rate constants for SΔTM D614G.

| SARS-CoV-2 spikes D614G | FRET-state occupancies (%) | | Rate constants ($sec^{-1}$) | |
|---|---|---|---|---|
| | Low-FRET (0.35) RBD "up"-conformation | High-FRET (0.65) RBD "down"-conformation | $k_1$ (Low to high FRET) | $k_{-1}$ (High to low FRET) |
| Unbound | 60.9±2.5 | 39.1±2.5 | 1.6±0.2 | 2.0±0.2 |
| + ACE2 | 74.5±2.2 | 25.5±2.2 | 0.7±0.1 | 2.0±0.4 |
| + Mab362IgG$_1$ | 57.4±1.1 | 42.6±1.1 | 2.0±0.1 | 2.6±0.1 |
| + Mab362IgA$_1$ | 62.1±1.2 | 37.9±1.2 | 1.6±0.1 | 2.6±0.1 |
| + REGN10987 | 64.1±1.9 | 35.9±1.9 | 2.0±0.2 | 3.7±0.3 |
| + S309 | 50.7±1.7 | 49.3±1.7 | 2.2±0.1 | 2.4±0.2 |
| + CR3022 | 60.0±2.6 | 40.0±2.6 | 1.1±0.2 | 1.8±0.2 |
| + 4A8 | 55.4±2.3 | 44.6±2.3 | 2.0±0.2 | 2.3±0.2 |
| + 1A9 | 64.7±1.7 | 35.3±1.7 | 1.7±0.1 | 3.2±0.2 |
| + 2G12 | 66.7±1.9 | 33.3±1.9 | 1.4±0.1 | 2.7±0.3 |

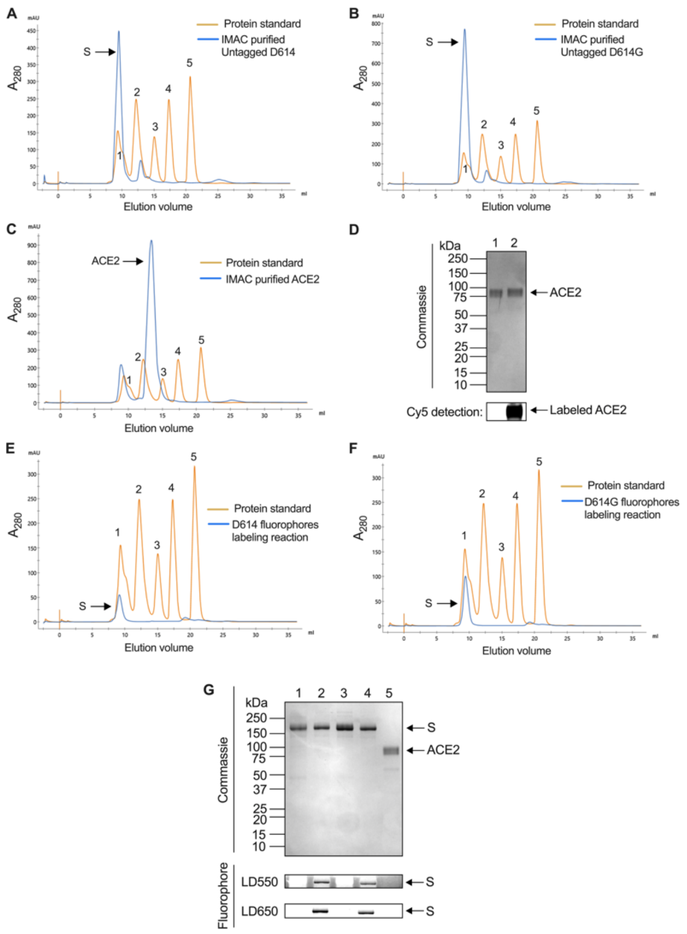


**Fig. S1. Purification and labeling of SARS-CoV-2 SΔTM and soluble ACE2.** (**A**) Chromatogram from size-exclusion chromatography (SEC) purification of SΔTM D614 (orange) as described in Material and Methods, with a gel filtration standard (Bio-Rad) overlaid. The 670, 158, 44, 17 and 1.35 kDa molecular weight references are numbered as 1, 2, 3, 4, and 5, respectively. (**B)** The same SEC data for SΔTM D614G, and (**C**) ACE2. (**D**) Cy5-labeling of purified ACE2 was analyzed by denaturing PAGE. Representative gel showing gel Coomassie staining (top) and Cy5 fluorescence (bottom) with unlabeled and labeled ACE2 in lanes 1 and 2, respectively. (**E**) SEC purification, displayed as in (A), for LD550/650-labeled A4-tagged SΔTM D614, and (**F**) SΔTM D614G hetero-trimers. (**G**) Denaturing PAGE analysis of purified unlabeled and labeled proteins. Labeling was performed as described in Materials and Methods. Coomassie staining (top) and in-gel fluorescence detection are as indicated. Lane 1: SΔTM D614; lane 2: LD550/650-labeled SΔTM D614; lane 3: SΔTM D614G; LD550/650-labeled SΔTM D614G; and, lane 5: unlabeled ACE2.

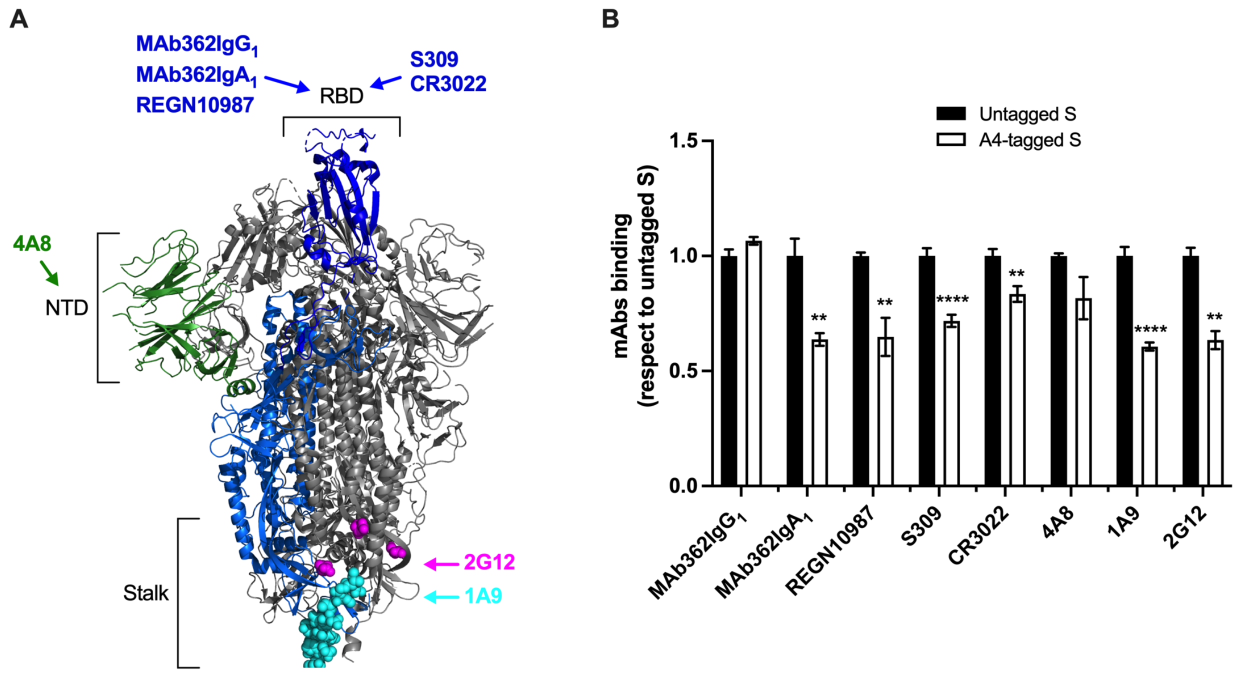


**Fig. S2. Characterization of SΔTM containing A4-peptide tags for site-specific fluorophore attachment.** (**A**) SARS-CoV-2 S domains and regions recognized by the mAbs used in this study are indicated. An S protomer with its RBD in the up conformation is depicted in colors: RBD, blue; NTD, dark green; cyan and magenta spheres, stalk epitopes for 1A9[48] and 2G12[49], respectively. The remainder of the protomer is shown in light blue. The remaining two protomers with their RBDs in the down conformation are shown in grey. Structure adapted from PDB 6VSB. (**B**) ELISA assay indicating mAb binding to A4-tagged SΔTM spikes as described in Material and Methods. Results from three independent experiments with two technical replicates each, using both untagged (white bars) or A4-tagged (black bars) SΔTM D614 spikes and the indicated mAbs are shown. Statistical significance was evaluated as described in Material and Methods and *P* values <0.05 were considered significant. Significance values are indicated as ***P*<0.01 and *****P*<0.0001.

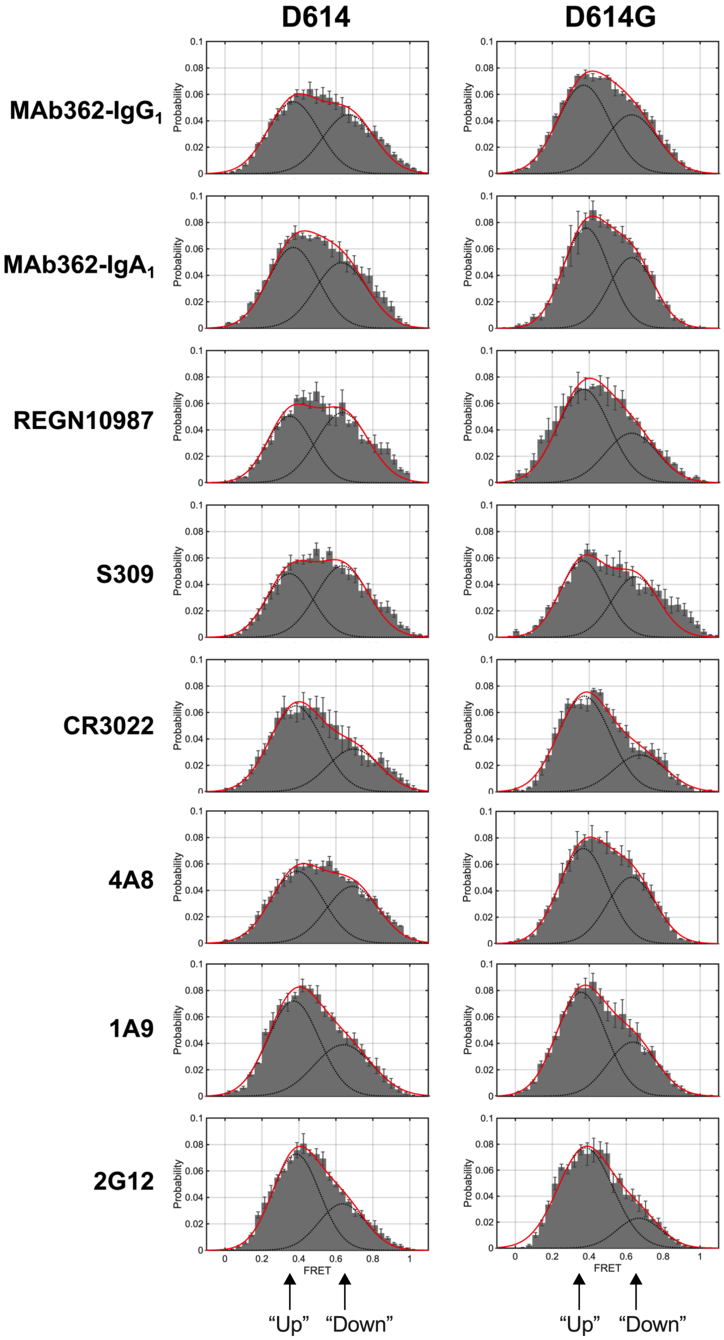


**Fig. S3. FRET histograms for SΔTM in the presence of mAbs.** Data are shown for both the D614 and D614G spikes in the presence of the indicated mAbs and are presented exactly as in Fig. 3A.
